# Muscle stem cell intramuscular delivery within hyaluronan methylcellulose improves engraftment efficiency and dispersion

**DOI:** 10.1101/245258

**Authors:** Sadegh Davoudi, Chih-Ying Chin, Michael C. Cooke, Roger Y. Tam, Molly S. Shoichet, Penney M. Gilbert

**Author notes:** Correspondence should be addressed to: Penney M. Gilbert, 164 College Street, Rosebrugh Building, Rm. 407, Toronto, Ontario M5S3E1.

## Abstract

Adult skeletal muscle tissue harbors the capacity for self-repair due to the presence of tissue resident muscle stem cells (MuSCs). Advances in the area of prospective MuSC isolation demonstrated the potential of cell transplantation therapy as a regenerative medicine strategy to restore strength and long-term regenerative capacity to aged, injured, or diseased skeletal muscle tissue. However, cell loss during ejection, limits to post-injection proliferation, and poor donor cell dispersion distal to the injection site are amongst hurdles to overcome to maximize MuSC transplant impact. Here, we assess a physical blend of hyaluronan and methylcellulose (HAMC) as a bioactive, shear thinning hydrogel cell delivery system to improve MuSC transplantation efficiency. Using in vivo transplantation studies, we found that the HAMC delivery system results in a >45% increase in the number of donor-derived fibers as compared to saline delivery. Furthermore, we observed a significant improvement in donor fiber dispersion when transplanted MuSCs were delivered in the HAMC hydrogel. Studies to assess primary myoblast and MuSC viability in HAMC culture revealed no differences compared to the media control even when the cells were first ejected through a syringe and needle or exposed to regenerating skeletal muscle extract to mimic the transplantation procedure. However, when we quantified absolute numbers, we found that more cells pass through the syringe and needle when delivered in HAMC. Culture in HAMC also increased the proportion of MuSCs in cell cycle, via a CD44-independent mechanism. An effect on myoblast proliferation was not observed, suggesting a hierarchical effect. Finally, a series of transplant studies indicated that HAMC delivery does not influence passive cell clearance or alter the host immune response, but instead may serve to support in vivo expansion by delaying differentiation following transplant. Therefore, we conclude that MuSC engraftment efficacy is improved by delivering the therapeutic cell population within HAMC.

## Introduction

Skeletal muscle is a striated muscle that facilitates voluntary movement, maintains posture, and aids in thermoregulation under the control of the nervous system^1,2^. A skeletal muscle tissue is comprised of multinucleated muscle fibers that are aligned with one another and organized into packed bundles to maximize contractile force. Amongst the mononucleated cells within a skeletal muscle are the ‘satellite cells’, named according to their anatomic positioning relative to the muscle fibers they reside atop^3^.

Satellite cells are a tissue resident stem cell required for the regenerative potential of healthy adult skeletal muscle tissue^3,4^. They express the paired box transcription factor Pax7 and are mitotically quiescent under homeostatic conditions^5–7^. Tissue insult activates satellite cells to divide and give rise to a population of transient amplifying cells co-expressing the myogenic regulatory factors MyoD and Myf5 along with Pax7. Eventually the transient amplifying population downregulates Pax7, MyoD, and Myf5 expression, and upregulates myogenin, thereby committing to exiting the cell cycle and fusing to reform the post-mitotic multinucleated muscle fibers^8–11^. In culture, the transient amplifying population undergoes a ‘crisis’ phase selecting for a subpopulation of cells referred to as primary myoblasts (pMBs), which can be passaged a limited number of times, and that are competent to fuse together into multinucleated muscle fibers upon mitogen withdrawal.

Myogenic cell transplantation is considered a putative treatment to restore localized strength and function to aged, injured, or diseased skeletal muscle. Early studies experimenting with pMB transplantation in clinical trials reported challenges including cell survival, rapid injection site clearance, poor donor cell dispersion, and limited contributions to tissue repair^12^. Within the last decade, methods to prospectively isolate mononucleated cells from murine^13–15^ and human^16–18^ skeletal muscle that possess muscle stem cell (MuSC) properties (i.e. self-renewal and differentiation) renewed hope for advocates of myogenic cell transplantation (reviewed in ^19^). Side-by-side comparisons of the in vivo expansion and regenerative potential of transplanted MuSCs compared to pMBs highlighted the therapeutic potency of MuSCs^13,20–24^. However, MuSCs are a relatively rare population of cells in skeletal muscle. Despite advances in the area of expanding MuSCs ex vivo while maintaining their regenerative capacity^17,25,26^, methods to produce clinically relevant numbers of this therapeutic cell population are still under development. Therefore, parallel efforts aimed at optimizing the transplantation procedure to maximize engraftment efficiency promise to synergize with MuSC ex vivo expansion studies to produce a clinically relevant therapy.

Synthetic and natural biomaterials are broadly studied in the context of putative skeletal regenerative medicine applications^24,27,28^. Biomaterials are an advantageous class of polymers due to the variety of biochemical and biophysical parameters that can be tuned to suit the regenerative medicine application. Surprisingly, the design of injectable biomaterials to improve MuSC transplantation efficiency following a simple intramuscular injection remains understudied despite the clear clinical value. In a recent study, peptide amphiphiles forming an injectable liquid crystalline scaffold were used to encapsulate and deliver murine muscle stem cells intramuscularly^29^. The peptide amphiphiles organized to form nanofibers and when extruded through a custom injection device into a tissue, the solution polymerizes via tissue resident divalent ions and the nanofibers align. Notably, MuSC engraftment efficiency was improved when delivered within the synthetic scaffold compared to the saline control. In vitro studies of the C2C12 immortalized cell line and mouse pMBs indicated that scaffold stiffness optimization efforts and the presence of aligned nanofiber maximized cell viability and proliferation, and may account for the observed in vivo benefits. To our knowledge, this was the first study evaluating an injectable cell delivery vehicle to improve MuSC transplantation efficiency.

In this study, we sought to further overcome translational challenges associated with MuSC intramuscular delivery through the study of a biomaterial scaffold shown to improve engraftment of other adult stem cell types^30–32^. HAMC is a hydrogel comprised of two components: hyaluronan (HA) and methylcellulose (MC)^33,34^. Hyaluronan is a natural polysaccharide found in all tissues including skeletal muscle extracellular matrix (ECM)^35,36^. Methylcellulose is a chemical compound derived from cellulose that, when dissolved in water, is liquid at low temperatures but forms a gel at higher temperatures^37^. HAMC is an injectable hydrogel due to its shear-thinning properties; while it is a liquid at lower temperatures, HAMC forms a gel at physiological temperatures^33^. HAMC is biodegradable and biocompatible and can attenuate the immune response in the brain and spinal cord^33,38^. Our previous studies demonstrated that using HAMC as a cell delivery vehicle for transplanting neural stem cells into the spinal cord^30^, retinal stem progenitor cells to the sub-retinal space^31^, retinal stem cell-derived rods to the retina^32^, and neural stem cells to the brain^32^, improves the survival, distribution, and contribution of the transplanted cells to regeneration.

HA is a naturally occurring ECM ligand^39^, and since myogenic cells express two of the known HA receptors (CD44 and RHAMM), it is reasonable to expect that HAMC exerts bioactive effects on encapsulated myogenic cells. For example, myoblasts express CD44^40^, which plays a role in regulating their migration, and differentiation^41^. Furthermore, studies of various cell types revealed that CD44-HA interactions promote cell growth and proliferation, and prevent apoptosis^42–46^.

In this study, we evaluate HAMC as a myogenic cell delivery vehicle, and investigate the cellular and molecular mechanisms by which it impacts the transplantation outcome. We report that intramuscular delivery within HAMC improves MuSC transplantation efficiency, without the need for a specialized delivery device, by increasing donor fiber numbers and their dispersion in the recipient tissue. From culture and in vivo studies, we conclude that the shear thinning property of HAMC increases the number of MuSCs that emerge during the ejection procedure, and that bioactive properties of HAMC delay MuSC differentiation, while promoting MuSC proliferation, and aiding in MuSC migration, thereby culminating in improved engraftment efficiency and dispersion compared to delivery in saline.

## Results

### Muscle stem cell delivery within HAMC improves engraftment efficiency and dispersion

To assess the influence of hyaluronin (HA) methylcellulose (MC) as a cell delivery vehicle for murine muscle stem cells (MuSCs), we performed transplantation assays in mice. Two days prior to transplant, we damaged the *tibialis anterior* (TA) muscles of recipient mice with a single intramuscular injection of BaCl_2_ to create a regenerative environment (Figure 1a). We prospectively isolated MuSCs from transgenic mice expressing GFP under the control of β-actin (Supplemental Figure 1a). The freshly isolated GFP^+^ MuSCs were suspended in saline or in 0.75:0.75% w/w HA:MC dissolved in saline and then transplanted intramuscularly into the injured TA muscle of immune-competent wild-type littermates to recapitulate a syngeneic MuSC transplantation therapy. To ensure that MuSCs were consistently transplanted into the center of the muscle, we added fluorescent microbeads into the cell injection mixture (Supplemental Figure 1b-c). 3-4 weeks after transplantation, we euthanized the animals and harvested the TA muscles (Figure 1a). The isolated TAs were sectioned and immunostained to visualize the contribution of GFP^+^ donor cells to the process of regeneration (Figure 1b).

**Figure 1.**
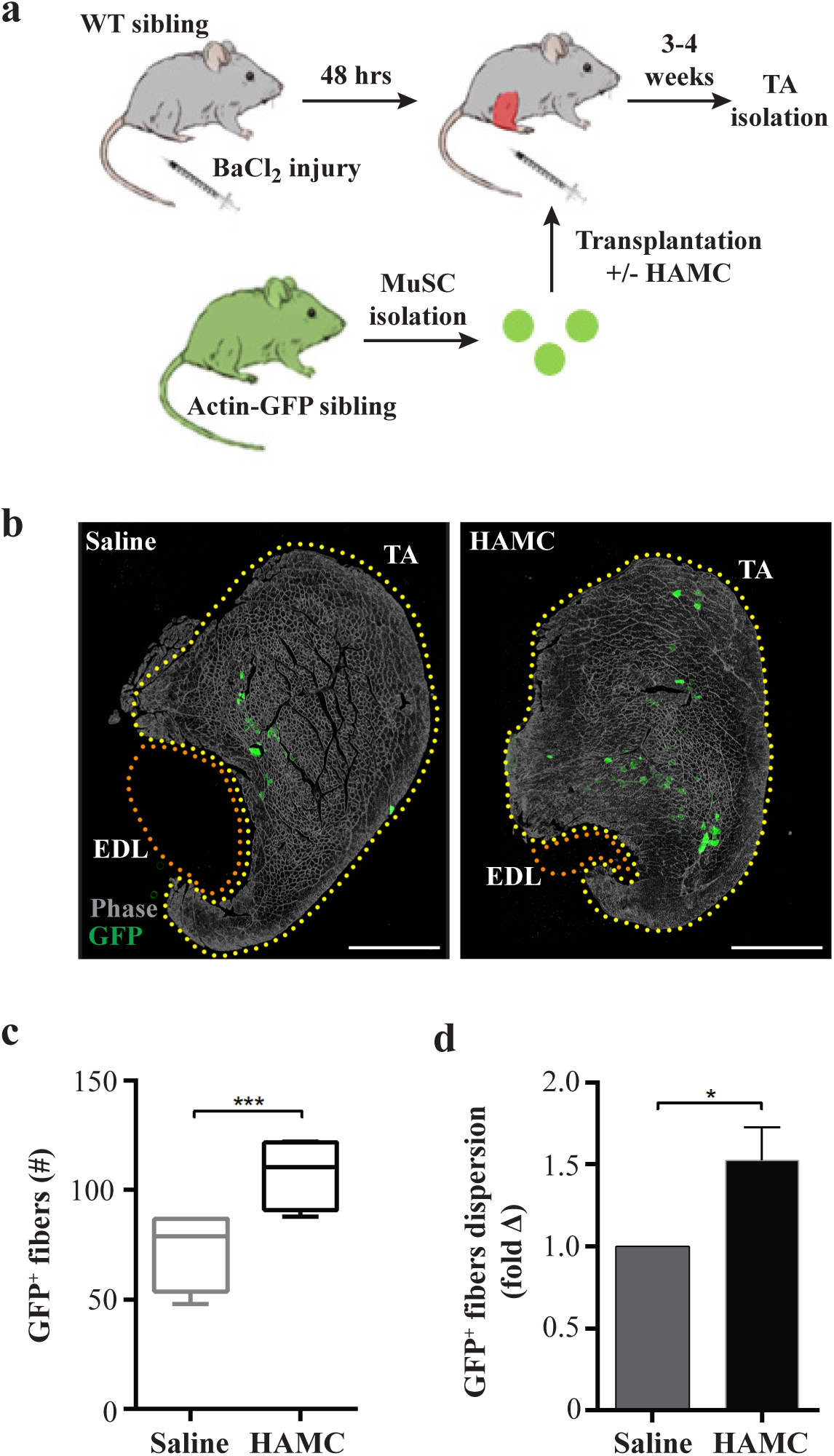
Delivery within HAMC improves muscle stem cell engraftment efficiency. **(a**) Schematic of muscle stem cell transplantation protocol. *Tibialis anterior* (TA) muscles of wild-type (WT) mice were injured with a single intramuscular injection of BaCl_2_. 48-hours following injury, muscle stem cells (MuSCs) freshly isolated from transgenic mice ubiquitously expressing Green Fluorescent Protein (GFP) were injected into the injured TA muscles of their WT siblings with (^+^) or without (−) hyaluronan methycellulose (HAMC). TA muscles were harvested 3-4 weeks post-transplantation and analyzed to determine the contribution of the transplanted MuSCs to muscle regeneration. **(b)** Representative tiled phase and epifluorescence composite images of TA and *extensor digitorum longus* (EDL) muscles immunostained for GFP at 4-weeks following transplantation with 10×10^3^ MuSCs delivered in saline control (left) or HAMC (right). Scale bar, 1 mm. **(c)** Graph portraying GFP^+^ fiber number following injection of 10×10^3^ MuSCs within saline (dark grey) or HAMC (black). n = 4. **(d)** Bar graph showing the fold change in the distribution of the GFP^+^ fibers (dispersion) 1-month following transplantation of 5×10^3^ or 10×10^3^ MuSCs within saline or HAMC control. n = 6. Error bars indicate SEM. Statistical significance determined by **(c)** paired or **(d)** unpaired student’s t-test where; p < 0.05.

We first performed a limiting dilution assay to assess the ability of transplanted GFP^+^ donor cells to outcompete endogenous wild-type MuSCs during the regeneration process in the immune-competent recipients. To this end, we transplanted different numbers of freshly isolated MuSCs within saline or HAMC and plotted the number of GFP positive fibers against the number of transplanted cells. We did not detect GFP^+^ donor fibers following transplantation of 1.5×10^3^ MuSCs, but observed a linear increase in GFP^+^ fibers following transplant of 5×10^3^ and 10×10^3^ GFP^+^ MuSCs (Supplemental Figure 2a) within saline (R^2^=0.81) or HAMC (R^2^=0.92). Interestingly, MuSC delivery within HAMC reduced transplant variability, an effect also noted in another study where MuSCs were delivered within a synthetic biomimetic scaffold^29^. Notably, we observed a >45% increase in the number of donor-derived (GFP^+^) fibers when MuSCs were delivered in HAMC compared to the saline control (Figure 1b-c and Supplemental Figure 2b). We observed no differences in donor fiber cross-sectional area (Supplemental Figure 2c-d), indicating that HAMC does not induce fiber hypertrophy. Interestingly, delivery within HAMC resulted in a significant increase in donor-derived muscle fiber dispersion (Figure 1b and d), suggesting that by optimizing the cell delivery vehicle it is possible to overcome a commonly observed MuSC transplant hurdle; failure of transplanted cells to engraft at sites distal to the injection site^47,48^. From these results, we conclude that transplanting MuSCs within HAMC improves on two stem cell therapy challenges: engraftment efficiency and cell dispersion.

### HAMC improves muscle stem cell ejection efficiency

Next, we sought to understand how HAMC influenced the transplanted cells using in vitro assays. Our aim was to recapitulate the complex tissue environment and the various conditions the transplanted cells are exposed to prior, during, and after the transplantation process, as well as during regeneration, to assess the effects of HAMC on early myogenic fate post-transplantation. We studied freshly isolated MuSCs and low passage primary myoblasts (pMBs) in culture to determine if HAMC elicits differential effects based on stem cell hierarchy status (i.e. quiescence, activation, transient amplifying).

HAMC was previously shown to improve cell viability^32^. Therefore, we tested the hypothesis that HAMC improves myogenic cell viability resulting in a greater number of therapeutic cells engrafting into the host. Hyaluronan is a natural component of the skeletal muscle microenvironment^39^ so we first assessed whether passive culture within bioactive HAMC impacts myogenic cell viability. We gently resuspended pMBs in HA:MC reconstituted in growth media using a wide-bore tip to reduce shear stress, and found no significant differences in the viability of the pMBs after 24 and 48 hours of culture as compared to the growth media control (Figures 2b-c).

**Figure 2.**
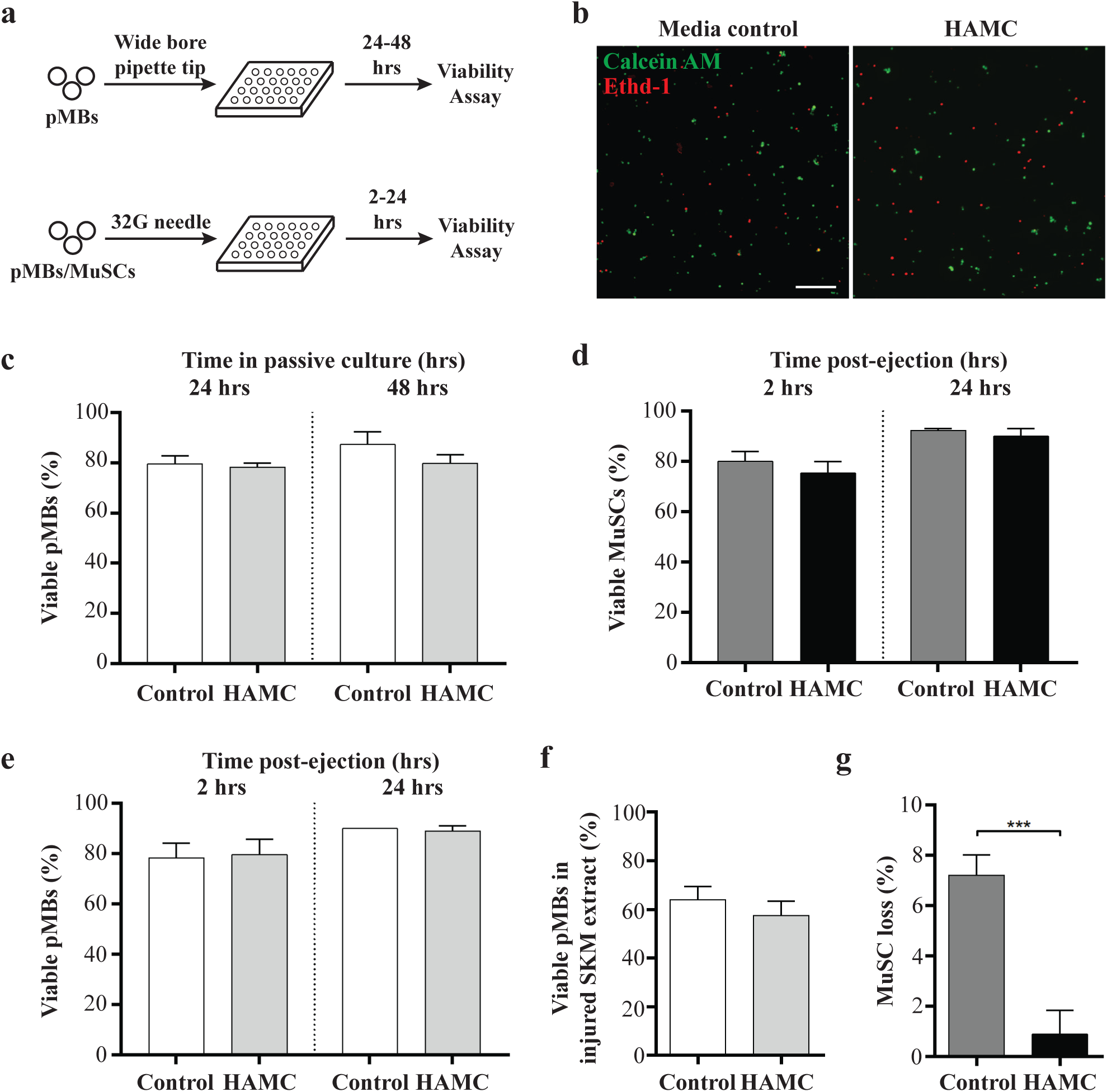
HAMC improves ejection efficiency without altering cultured myogenic cell viability. **(a)** Schematic of cell viability assays comparing HAMC to media control. Primary myoblasts (pMBs) and muscle stem cells (MuSCs) were plated with or without HAMC using a wide-bore pipette tip (top) or a syringe with a 32G needle (bottom) into a 96 well plate. At the time-points indicated, the calcein AM and ethidium homodimer ‘live/dead™ assay’ was used to assess cell viability in each condition. **(b)** Representative epifluorescence images of pMBs cultured in control media or HAMC and co-stained with calcein AM (green, live) and ethidium homodimer (Ethd-1, red, dead). Scale bar, 200 µm. **(c)** Bar graph indicating the viability of pMBs 24 and 48 hours (hrs) after plating in media control (dark grey) compared to HAMC (black) culture. n = 3. **(d-e)** Bar graphs showing the viability of (**d**) MuSCs and (**e**) pMBs 2 and 24 hours following ejection through a syringe equipped with a 32G needle when delivered within media (MuSCs, dark grey; pMBs, white) or HAMC (MuSCs, black; pMBs, light grey). n = 3. **(f)** Bar graph indicating the viability of pMBs 24 hours after ejection through a syringe and 32G needle when delivered within media (white) or HAMC (light grey) into a 1mg / mL solution of skeletal muscle (SKM) extract that was prepared 48 hours after a BaCl_2_-induced tissue injury. n = 3. **(g)** Bar graph showing the percentage of MuSCs lost during ejection through a syringe and 32G needle when delivered within media (dark grey) as compared to HAMC (black). n = 3. Error bars indicate SEM. Statistical significance determined by student’s t-test where; p < 0.05.

HAMC is a shear thinning hydrogel^33^, so we next tested whether the reduced shear stress during ejection through a narrow needle protected the cells from damage during the transplant process (Figure 2a). Here we quantified the proportion of viable MuSCs (Figure 2d) and pMBs (Figure 2e) in the culture well 2 hours or 24 hours after ejection through a 10 µL Hamilton syringe equipped with a 32G needle. Again, we observed no differences in cell viability when comparing myogenic cell delivery in HAMC versus the culture media control (Figures 2d-e). Our results are in line with prior pMB studies concluding that passage through a syringe and needle partially damages the cell membrane, but does not result in cell death in vitro^49–51^.

If the ejection process induces membrane damage, we posited that exposure to biomolecules present in the hostile regenerative environment might push the transplanted cells towards death, and that perhaps HAMC protects against this. To simulate the regenerating environment of injured skeletal muscle that the donor cells are injected into, we injured the TA of C57Bl6/N mice, harvested the injured tissue 48 hours later, and prepared a tissue extract^52^. We then resuspended primary myoblasts in HAMC or media control and ejected the cell suspensions through a Hamilton syringe equipped with a 31G needle directly into a culture well containing the injured skeletal muscle tissue extract. As expected, ejection and culture in injured tissue extract resulted in an overall reduction in cell viability (Figure 2f; ~60%) compared to the growth media (Figure 2c and e; ~80%), and delivery in HAMC did not afford protection from death in this setting (Figure 2f). These results suggest that HAMC does not improve the viability of myogenic cells that pass through the syringe and needle.

All of our viability studies to this point focused on analyzing the cells after ejection through the syringe and needle. Interestingly, when we quantified the total number of MuSCs that passed through the syringe and needle into the culture dish, we observed a greater number of cells when delivered in HAMC compared to the saline control (Figure 2g). This translated to a 6% reduction in stem cell loss during the ejection procedure. Therefore, we conclude that myogenic cell delivery in HAMC does not modify the viability of the cells that emerge from the needle. However, HAMC ultimately increases ejection efficiency by protecting cells from obliteration during the ejection or by preventing cells from becoming lodged in the syringe or needle during ejection, thereby resulting in an increase in the total number of myogenic cells transplanted into the tissue.

### HAMC promotes MuSC proliferation via a CD44-independent mechanism

Based on our limiting dilution analysis (Supplemental Figure 2a), the modest improvement in ejection efficiency (6%; Figure 2f) does not fully account for the >45% increase in donor derived fibers observed when MuSCs are delivered in HAMC (Figure 1c). Therefore, we investigated the possibility that HAMC may also influence myogenic cell proliferation. Freshly isolated MuSCs were cultured in growth media or HA:MC reconstituted in growth media for a period of 72 hours and pulsed with 5-ethynyl-2’-deoxyuridine (EdU) during the final 12 hours of culture (Figure 3a). Intriguingly, we found that the number of cells that incorporated EdU was ~14% higher when embedded in HAMC compared to culture in the growth media control condition (Figure 3b, left). In addition to revealing another benefit of using HAMC for MuSC delivery, these results suggest that HAMC either pushes a greater proportion of MuSCs to activate from quiescence or it increases the proliferation rate of MuSCs following activation.

**Figure 3.**
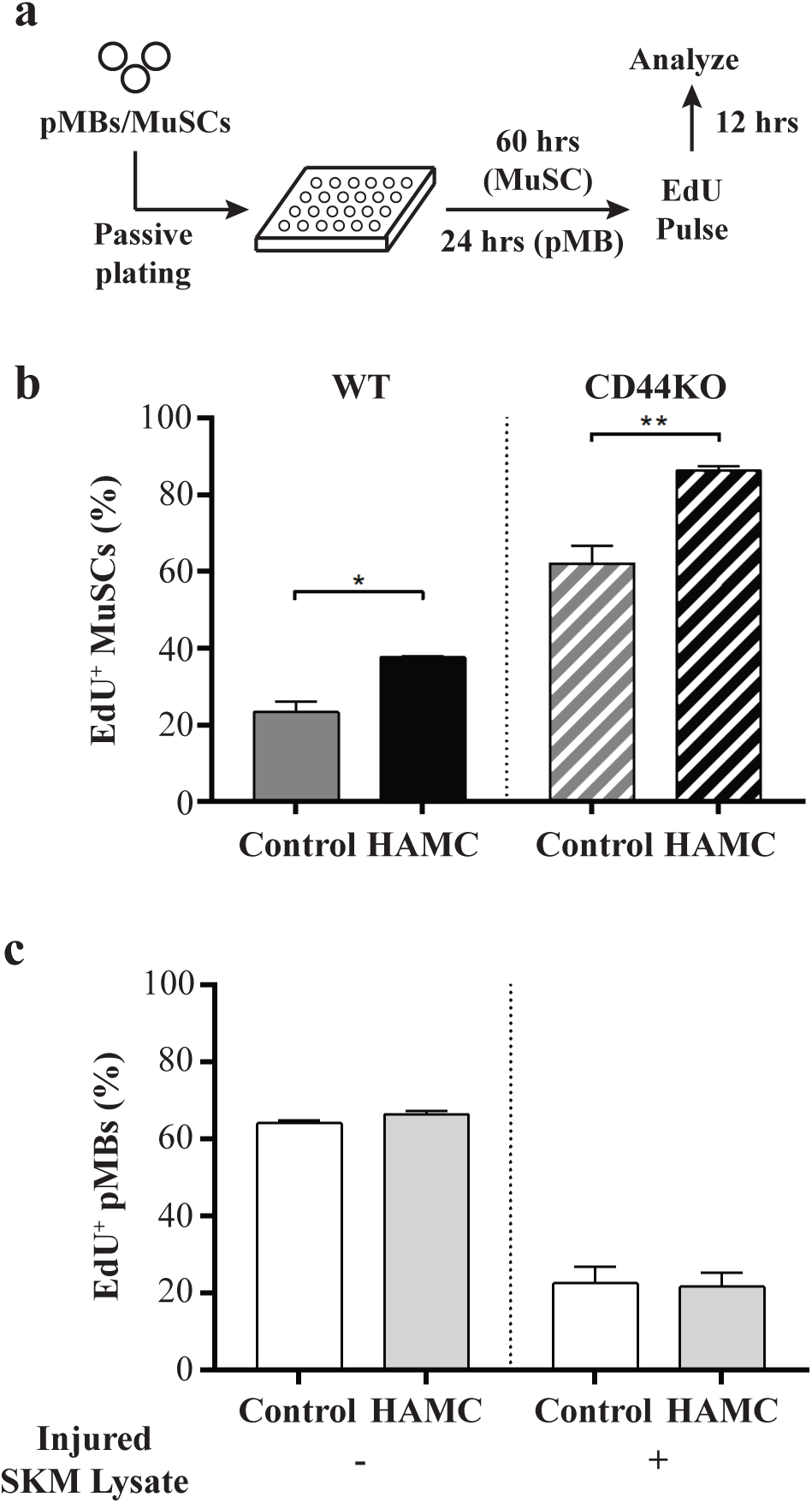
HAMC influences MuSC proliferation via a CD44-independent mechanism. **(a)** Schematic of 5-ethynyl-2’-deoxyuridine (EdU) assay comparing myogenic cell proliferation in media control as compared to HAMC culture. Primary myoblasts (pMBs) or muscle stem cells (MuSCs) were plated into a 96 well plate with or without HAMC using a wide-bore pipette tip. At the culture time-points indicated, EdU was added to the culture media for 12 hours (hrs). Cells were then fixed, stained, and analyzed for EdU incorporation. **(b)** Bar graphs showing the percentage of MuSCs freshly isolated from **(left)** wild-type (WT; n=6) or **(right)** CD44 knock-out (CD44KO; n=3) mice that incorporate EdU after 72 hours of culture in media control (WT, dark grey; CD44KO, patterned dark grey) or HAMC (WT, black; CD44KO, patterned black) when pulsed with EdU for 12 hours prior to the analysis time-point. **(c)** Bar graphs depicting the percentage of primary myoblasts that incorporated EdU after 24 hours of culture when plated in growth media control (white) or HAMC (light grey) in the presence (**right**) or absence (**left**) of injured skeletal muscle (SKM) lysate when pulsed with EdU for 12 hours prior to the analysis time-point. n=3. Error bars indicate SEM. Statistical significance determined by student’s t-test where; p < 0.05.

In the first week of muscle repair, MuSCs activate and proliferate to give rise to a transient amplifying progenitor pool that will ultimately fuse to produce nascent muscle fibers on the third day of repair in the regenerating tissue. Since prior studies showed that HAMC remains at the injection site for as many as 7 days^31^, we next investigated the effects of HAMC on pMB (i.e. transient amplifying progenitors) cell cycle entry. pMBs were cultured in growth media or HAMC reconstituted in growth media for a period of 36 hours and pulsed with EdU during the final 12 hours of culture. Interestingly, similar proportions of pMBs incorporated EdU incorporation when cultured in HAMC compared to the growth media control (Figure 3c, left). We then investigated the effect HAMC on pMB cell cycle entry in the context of the regenerating environment by ejecting cells into injured skeletal muscle tissue extract. In this context, we observed an overall lower incidence of EdU incorporation, with no significant differences observed between the HAMC and control conditions (Figure 3c, right). Together, these data (Figure 3b-c) suggest that HAMC-induced effects on myogenic cell cycle entry are limited to MuSCs, and not their downstream progeny, revealing an intriguing hierarchical bias.

Signaling through the HA-CD44 receptor-ligand axis positively regulates cell cycle entry in a wide variety of cell types^42–46^. Therefore, we sought to determine whether the proliferative influence of HAMC on MuSCs was mediated by CD44. First, we used flow cytometry to determine if myogenic cells express and present the CD44 receptor. Consistent with recent reports^53^, we found that CD44 is not expressed on freshly isolated MuSCs, while activated MuSCs and primary myoblasts both have CD44 cell surface expression (Supplemental Figure 3). Next, freshly isolated MuSCs from CD44^−/−^ mice were cultured in HAMC or the growth media control for 72 hours and pulsed with EdU for the final 12 hours of culture (Figure 3a). In contrast with reports for other cell types^32,42,43,46,54^, loss of CD44 elicited a dramatic increase in the proportion of MuSCs entering cell cycle compared to the wild-type control (Figure 3b; grey solid and patterned bars). Furthermore, similar to our wild-type MuSC study (Figure 3b; left), culture within HAMC increased the proportion of CD44KO MuSCs that incorporated EdU during the final 12 hours of the 3-day culture period (Figure 3b; right). Taken together, our results suggest that HAMC culture leads to an increase in MuSC, and not pMB proliferation, via a CD44-independent mechanism.

### HAMC does not modify the skeletal muscle innate immune response

One of the common causes of cell death post-transplantation is the inflammatory reaction, with the neutrophil response inducing particularly deleterious effects on cell survival^55,56^. Prior studies showed that HAMC attenuates the inflammatory response (i.e. the presence of microglia and astrocytes) in the brain and spinal cord^33,38^. However, the effect of HAMC on the inflammatory environment in skeletal muscle has not been investigated. To narrow in on additional mechanisms by which HAMC improves the engraftment efficiency of the transplanted cells, we investigated the presence of neutrophils and macrophages following HAMC injection. First, we injected a barium chloride solution intramuscularly into the TA muscles of C57Bl/6N wild-type mice. 48 hours later we injected saline +/− HAMC (and mixed with fluorescent beads) intramuscularly into the center of the regenerating TA muscle group. At 2 and 24 hours post-injection, we harvested the tissues for immunohistological analysis (Figure 4a).

**Figure 4.**
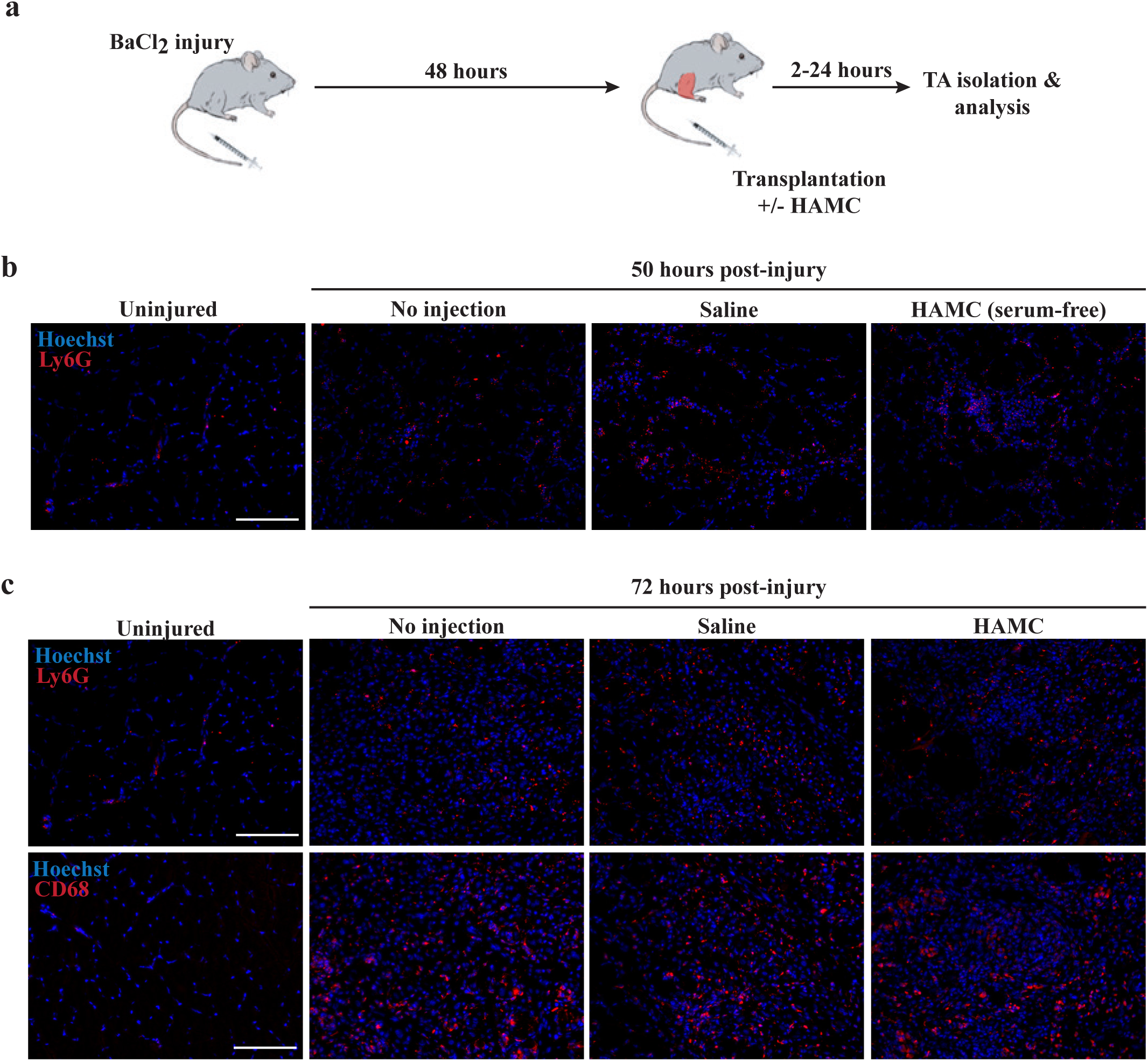
HAMC does not modify Ly6G^+^ or CD68^+^ immune cell incidence in the first 24 hours post-transplantation. **(a)** Schematic of experiments designed to assess the short-term effects of HAMC transplantation on the early regeneration immune response. *Tibialis anterior* (TA) muscles of wild-type (WT) mice were injured with BaCl_2_. 48 hours post-injury, HAMC hydrogel or a control media was injected into injured TA muscles of WT mice. The TA muscles were isolated 2 or 24 hours post-transplantation (equivalent to 50 and 72 hours post-injury) and analyzed immunohistologically to visualize the immune response at early time points post-injection. **(b)** Representative epifluorescence images of neutrophil (Ly6G, red) distribution in uninjured skeletal muscle (far left) and injured tissue 50 hours post BaCl_2_-injury (middle left). Images of neutrophil presence 2 hours post-injection of saline control (middle right) or HAMC (far right) into the pre-injured TA are also shown. Scale bar, 100 µm. **(c)** Representative epifluorescence images of neutrophil (top, Ly6G) and macrophage (bottom, CD68) distribution in uninjured (far left) and injured tissue 72 hours post BaCl2-injury (middle left). Images of neutrophil and macrophage presence 24 hours post-injection of saline control (middle right) or HAMC (far right) into the pre-injured TA are also shown. Scale bar, 100 µm.

As expected, neutrophils and macrophages of the innate immune system, expressing Ly6G and CD68, respectively, have a scarce presence in healthy, uninjured TA muscle (Figure 4b-c, far left panels). However, their incidence dramatically increased 50 hours (Figure 4b; middle left panel) and 72 hours (Figure 4c; middle left panel) after a barium chloride-mediated tissue injury. HAMC injection (+/− serum) did not induce gross alterations in the incidence or localization of neutrophils 50 hours after injury (Figure 4b and Supplemental Figure 4) or the neutrophils (Figure 4c; top panels) and macrophages (Figure 4c; bottom panels) 72 hours (Figure 4c and Supplemental Figure 5) after injury, when compared to saline control injections. Interestingly, the presence of bovine serum in the injection media did not alter the results at the time-points tested (Supplemental Figures 4 & 5). Based on these results, we conclude that HAMC addition does not modify days 2 – 3 of the skeletal muscle regeneration innate immune response. However, it remains to be determined whether HAMC influences earlier time-points in the repair process or whether HAMC influences immune cell behavior and / or secretome.

### HAMC prevents the active clearance of transplanted myogenic cells

Previous studies revealed that a majority of transplanted myoblasts do not withstand the first 4 days post-transplantation^56^. Loss of cells over these early time-points might be due to death, but might also be owed to passive clearance or active (e.g. immune cell mediated) clearance mechanisms. Therefore, we designed a series of experiments to determine whether HAMC prevents cell clearance post-transplantation. Since HAMC did not influence pMB proliferation in culture (Figure 3c), we utilized pMBs in these studies to avoid proliferation as a confounding parameter.

We transplanted 1×10^5^ GFP^+^ pMBs into the TA of WT littermate control animals 48 hours after a BaCl_2_-induced injury. pMBs were delivered within saline or HAMC and after 2, 24, or 48 hours, we harvested and dissociated the TA into a mononucleated cell slurry. With flow cytometry, we quantified the number of retrieved cells (Figure 5a-b). No significant differences were observed in the number of GFP^+^ cells retrieved 2 hours after transplanting GFP^+^ pMBs in HAMC versus saline (Figure 5c-d), suggesting that the HAMC hydrogel did not serve to prevent the passive clearance of the injected cell population. At later time-points post-injection, we expect cell clearance is due to active, immune cell-mediated mechanisms. No differences were observed 24 hours post-injection (Figure 5c,e), but at 48 hours we observed a ~1.5-fold increase in the number of cells retrieved when they were delivered in HAMC (Figure 5c,f). Since we observed no differences in cell retrieval at 2 and 24 hours post-transplant, we conclude that HAMC does not influence passive or active cell clearance. However, since the 48-hour post-transplant analysis time-point coincides with the period when mononucleated cells fuse to form multinucleated muscle fibers during regeneration, we conclude that HAMC may serve to delay MuSC differentiation and instead preserves an extended proliferation window to augment transplanted MuSC engraftment efficiency (Figure 5f).

**Figure 5.**
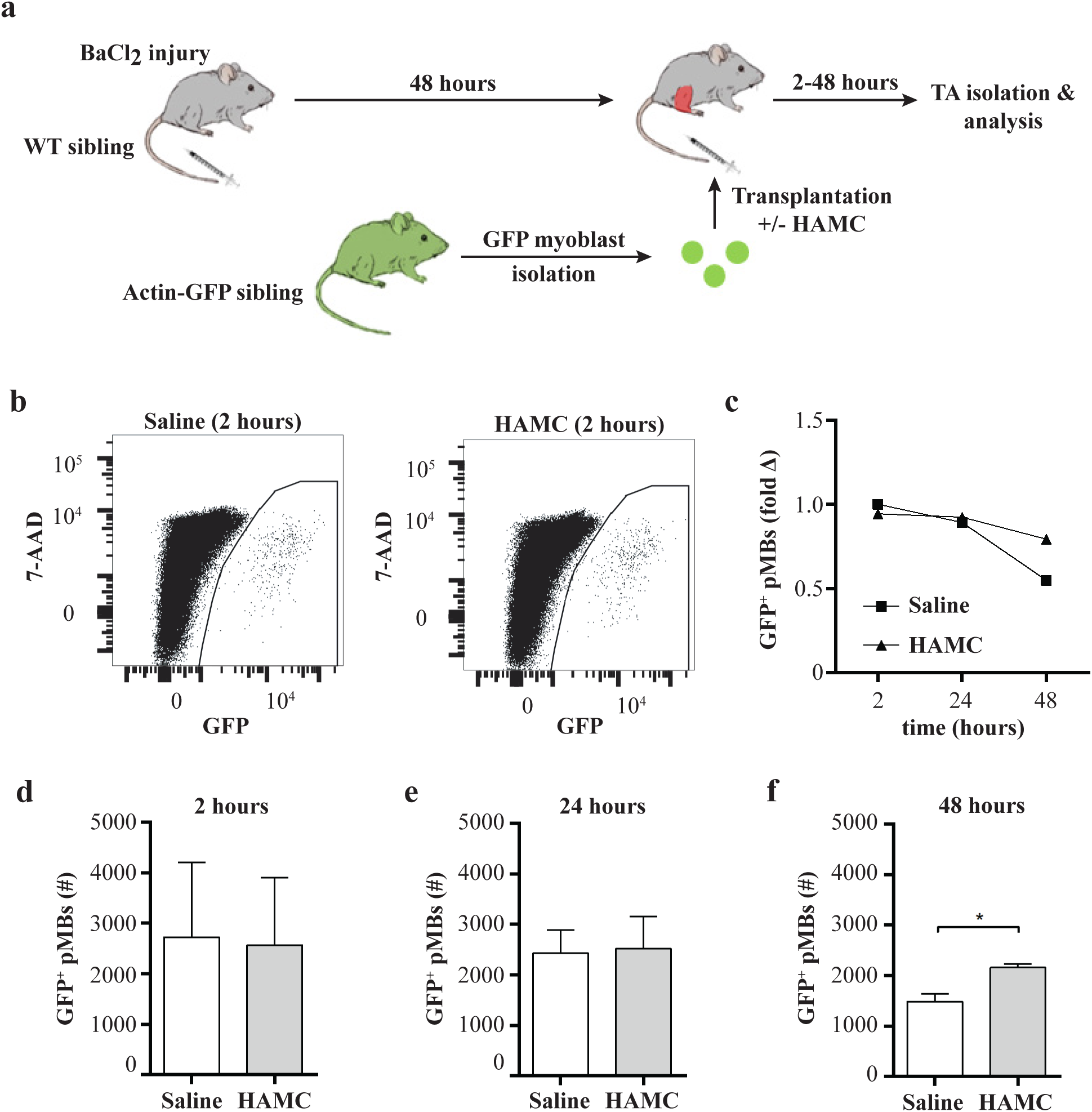
HAMC increases retention of transplanted cells post-transplantation. **(a)** Schematic of experiments aimed at determining the effect of HAMC on mononucleated cell retention within the first two days post-transplant. *Tibialis anterior* (TA) muscles of wild-type (WT) mice were injured with BaCl_2_. 48 hours post-injury, 10^5^ actin-GFP primary myoblasts were transplanted intramuscularly into the injured TA muscles within saline or HAMC hydrogel. The TA muscles were isolated 2, 24, or 48 hours post-transplantation, enzymatically digested, and analyzed by flow cytometry to quantify the number of GFP^+^ mononucleated cells. **(b)** Representative flow plots of retrieved GFP^+^ myoblasts 2 hours post-transplantation within saline (left) or HAMC (right). **(c)** Scatter plot displaying the fold change in GFP^+^ myoblasts retrieved over time when delivered within HAMC (triangle) compared to saline (square) where data are normalized to the number of cells retrieved at 2 hours post-transplantation within saline. **(d-f)** Bar graphs replotting the raw data from **(c)** to indicate the absolute number of GFP^+^ myoblasts retrieved from enzymatically digested skeletal muscle tissue (**d**; n = 3) 2 hours, (**e**; n=7) 24 hours, and (**f**; n = 3) 48 hours after intramuscular delivery within HAMC (light grey) compared to saline control (white). Error bars indicate SEM. Statistical significance determined by student’s t-test where; p < 0.05.

Based on our culture and in vivo results, we conclude that HAMC improves the engraftment efficiency of transplanted MuSCs by supporting the delivery of an overall greater number of cells, by promoting proliferation, and by delaying differentiation, which in turn serves to augment in vivo expansion. Furthermore, by visualizing the position of the fluorescent beads that were co-delivered with the MuSCs relative to engrafted GFP^+^ muscle fibers (Supplemental Figure 1b,c), it is clear that passive cell dissemination does not account for the HAMC-mediated improvement in donor fiber dispersion. Since, CD44 is also known to promote myogenic cell migration, it is enticing to speculate that improved cell dispersion is due to the active engagement of CD44 receptors mediated by HA-induced CD44 receptor ligation.

## Discussion

Our results show that the application of HAMC, a shear thinning hydrogel cell delivery vehicle, to MuSC intramuscular transplantation, increases engraftment efficacy and dispersion. Our mechanistic studies suggest that enhanced engraftment effects are owed to both HAMC material and bioactive properties. To our knowledge, this is the first report of a bioactive delivery vehicle improving MuSC engraftment efficiency and the first delivery vehicle of any kind augmenting MuSC post-transplant dispersion.

We sought to explore MuSC transplantation efficacy in the context of an intact immune response since the immune system is known to contribute to the process of skeletal muscle repair (reviewed in ^57^). In contrast, most studies in the area of MuSC transplantation utilize immuno-compromised and hindlimb irradiated animals as recipients. As such, we first performed limiting dilution analyses to ensure that the number of GFP^+^ MuSCs transplanted into syngeneic immune competent sibling recipients was in the linear range of engraftment for our saline and HAMC delivery conditions. Plotting the number of histologically detected GFP^+^ fibers versus the number of transplanted cells revealed that even at our highest transplant number (1×10^4^), we were still within the linear range, as we had not yet reached a fiber engraftment plateau (Supplemental Fig. 2a)^19,58–60^. Since the transplanted cells compete with endogenous MuSCs, this experimental model and data may be useful to future studies aimed at recapitulating autologous MuSC transfer.

We found that delivering MuSCs intramuscularly within HAMC rather than within saline resulted in an overall greater number of GFP^+^ donor-derived fibers in transplant recipients (Figure 1a-c and Supplemental Figure 2a-b). While HAMC did not induce fiber hypertrophy that was detectable after 4-weeks of regeneration (Supplemental Figure 2c-d), we cannot rule out the possibility that HAMC may influence muscle fiber hypertrophy at earlier time-points of repair. However, given that skeletal muscle contains copious amounts of hyaluronic acid, it seems unlikely that HAMC would exert an additive effect in this regard.

To identify a cellular mechanism to understand the positive impact of HAMC on MuSC engraftment efficacy, we evaluated several culture scenarios to explore the hypothesis that HAMC may improve transplanted MuSC viability. By simply mixing pMBs in HAMC and quantifying cell viability after 24 and 48 hours of exposure, we detected no differences between the two culture conditions (Figure 2a-c). Shear thinning hydrogels can improve cell transplantation outcome by reducing the extensional flow at the entrance of the needle, which otherwise will induce membrane damage during the ejection^61^. However, when we assessed the viability of MuSCs or pMBs that emerged from the syringe and needle when encapsulated in HAMC, we observed no differences compared with ejection within saline control (Figure 2d-e). This result aligns with that of previous studies^49–51^ showing that pMBs that survive the ejection process maintain high viability in culture. Since it was previously reported that the ejection process compromises membrane integrity^49^, we tested the hypothesis that HAMC may protect myogenic cells from death signals they encounter in the hostile transplant environment. While the viability of pMBs ejected into an injured skeletal muscle extract was decreased compared to ejection into regular growth media (Figure 2e-f), HAMC did not elicit a protective effect in this context either (Figure 2f). We note that our results with myogenic cells conflict with our prior studies demonstrating that HAMC encapsulation increases the proportion of viable cells after incubation^31,32,34,62^, and we expect this is an example of a HAMC cell type specific effect.

Finally, we evaluated the total number of MuSCs that emerged from the syringe and needle when delivered in HAMC compared to the saline control. In the highly controlled setting of culture, we found that more MuSCs emerged from the syringe and needle when delivered in HAMC (~6%). We expect that HAMC either protects MuSCs from being obliterated during ejection or, since hyaluronan is made of polysaccharides, the sugar groups may act as a cellular ‘slip-n-slide’ to prevent cells from becoming lodged in the syringe or needle. Interestingly, when we harvested and dissociated TA muscles to retrieve and quantify myogenic cells shortly after intramuscular injection (2 hours), we saw no difference in the total number of cells retrieved when delivered in HAMC compared to saline (Figure 5a-d). However, experimental error introduced during the process of tissue digestion, mononucleate cell retrieval, and flow cytometric analysis may introduce error that effectively masks the 6% difference in transplanted cell number.

Ultimately a 6% increase in delivered MuSCs does not fully account for the 1.5-fold increase in donor derived fibers that we observed when using HAMC as the delivery vehicle. As such, we explored the possibility that HAMC encapsulation promotes MuSC proliferation potential. Indeed, we observed a greater proportion of HAMC-encapsulated MuSCs in cell cycle (~14%) when we assessed EdU incorporation in the freshly isolated MuSC population after an EdU pulse in the final 12 hours of the 72 hour culture period (Figure 3a-b). Interestingly, this effect appears to be specific to activated MuSCs since pMB cell cycle status was insensitive to the presence of HAMC (Figure 3a,c). Even applying an additional environmental pressure, such as exposure to lysate from regenerating skeletal muscle, did not uncover HAMC-mediated effects on pMB proliferation (Figure 3a,c), thereby uncovering a MuSC-specific effect of HAMC.

HA-CD44 interactions induce ERK1/2 and PI3K-AKT signaling pathway activation, leading to the growth and proliferation of other cell types^36,43,44,63,64^. Consistently, inhibition of CD44-HA results in the inhibition of growth^46^ and cell apoptosis^45^ of other cell types. Since we previously showed that HAMC-mediated effects on cell proliferation were mediated by CD44^32,42,43,46,54^, we sought to determine whether the CD44-HA signaling axis was an underlying mechanism of HAMC-mediated MuSC proliferation. We first assessed cell surface expression of CD44 on myogenic cells (Supplemental Figure 3) and found that activated MuSCs and pMBs uniformly express the receptor, but, similar to recent studies using Cytof^53^, only a small subpopulation of freshly isolated MuSCs presented the receptor. When we prospectively isolated MuSCs from CD44^−/−^ mice, encapsulated and cultured the MuSCs in HAMC for 72 hours, and then assessed EdU incorporation, we still observed a greater proportion of MuSCs in cell cycle compared to the control condition (Figure 3a,b). Interestingly, comparing the control conditions (Figure 3b grey solid and striped bars), revealed a greater proportion of EdU incorporated MuSCs in the absence of the CD44 receptor at this time-point. It is possible that loss of CD44 in MuSCs releases a brake on MuSC proliferation. However, given that a prior study noted a delay in differentiation when studying CD44^−/−^ pMBs in culture^41^, our MuSC results may instead reflect delayed activation. Regardless, the HAMC-mediated boost in MuSC proliferation cannot be accounted for by a CD44-HA interaction, and instead implicate a role for another HA receptor such as Receptor for Hyaluronan-Mediated-Motility (RHAMM).

Since HAMC gels at physiological temperatures, we hypothesized that delivery in HAMC might protect MuSCs from being passively cleared in the earliest time-points after injection. However, when we collected GFP^+^ cells 2 hours after intramuscular injection, flow cytometric analysis revealed no difference in the total number of retrieved cells when delivered in HAMC compared to saline.

Next, we designed a series of experiments to determine whether HAMC might modify the skeletal muscle immune environment to support the observed improvements in MuSC engraftment efficiency. High cell death, especially during the first 4 days post-transplantation, is a common reason for transplantation failure^19^. The skeletal muscle inflammatory response after exercise or acute injury includes the invasion of neutrophils, macrophages, and natural killer (NK) cells^56,57^. Studies showed that neutrophils secrete proteases to break down cellular debris, and in the process, can cause damage and death to bystander healthy cells through release of cytokines and cytotoxins^55,57,65^. Macrophages and NK cells, on the other hand, do not appear to be detrimental to the fate of transplanted cells, and are more targeted in their activities^55,66^. Since previous studies on HAMC showed it to attenuate inflammation in the brain and spinal cord by reducing the presence of astrocytes and microglia, respectively^33,38^, we next assessed whether HAMC elicits similar effects in skeletal muscle. In this regard, we mimicked our transplantation regime by injuring immune-competent C57Bl/6J hindlimbs with a single injection of BaCl_2_ and then 2 days later, injected saline or HAMC, and then harvested the tissue for histological analysis 2 hours or 24 hours later (Figure 4 and Supplemental Figures 4-5). We observed no gross differences in the distribution or accumulation of Ly6G^+^ neutrophil and CD68^+^ macrophage cell populations compared to uninjected and saline injected controls.

Our histological analysis does not out-rule the possibility that perhaps HAMC shields the transplanted cells from deleterious interactions with the immune system that might then manifest as an increase in the total number of transplanted cells present over time. Indeed, previous studies showed that blocking interactions between neutrophils and transplanted cells increases their chance of survival^50^. Additionally, treating cells with fibronectin and vitronectin increased the survival of transplanted myoblasts, and reduced anoikis^56,67^. Therefore, the HA in HAMC might prevent anoikis by activating pathways involved in cell survival and growth such as ERK1 or PI3K-AKT pathways^36^. To test this theory, we transplanted GFP^+^ pMBs into pre-injured tissue and then retrieved and enumerated the cells using flow cytometry 24 hours and 48 hours later (Figure 5c,e,f). Since our culture studies concluded that HAMC does not influence pMB proliferation, we used pMBs in our retrieval studies to avoid the confounding influence of proliferation on cell count studies. While no differences in retrieved cell number were uncovered 24 hours after injection, at 48 hours a statistically significant decrease in total retrieved cells was observed in the saline control condition while the HAMC condition seemed to maintain a constant number of cells when assessed at 2, 24, and 48 hours post-transplant (Figure 5a-f). This data supports a protective effect of HAMC in preventing cell death and/or clearance by immune cells. However, 4 days after injury (i.e. 48 hours after transplant in our experimental regime; Figure 5a) corresponds to the time-point when myogenic progenitors fuse to form multinucleated muscle fibers. Therefore, we favor the possibility that the reduction in the number of mononucleated cells retrieved at 48 hours can be explained by fusion and that HAMC serves to delay differentiation.

In addition to delivering a greater number of cells and influencing proliferation, our HAMC data indicate that the delivery scaffold improves the migration of transplanted cell resulting in greater dispersion of donor-derived muscle fibers (Figure 1d). Studies showed that the interaction between HA and its receptors, including CD44 and RHAMM, leads to increased cell motility, and is involved in supporting the migration of many cell types including pMBs^41,68–70^. Additionally, a recent study suggested that a biphasic relationship exists between CD44 expression and survival and migratory behavior of glial cells in that increased CD44 expression resulted in faster migration rates and lower cell survival whereas lower CD44 expression lead to higher survival and less migration^71^. Hence, the increased dispersion of GFP^+^ donor-derived fibers could be a result of the increased HA presence in the local milieu^72^ or HA receptor ligation induced in the process of MuSC encapsulation and delivery. Since pretreating MuSCs with recombinant Wnt7a prior to transplant also influences cellular dispersion^73^, it would be interesting to determine whether HAMC delivery affords an additive effect on this engraftment metric.

Ultimately, an important end goal of MuSC transplantation is improved skeletal muscle function. It is unclear whether the 1.5-fold increase in donor derived fibers reported in our study translates to an increase in muscle force generation^73^. However, because the transplantation benefits we observed were largely attributed to bioactive influences of HAMC, we predict that modifying the recently reported peptide amphiphile-based MuSC delivery scaffold to include hyaluronan peptide sequences would serve to boost the efficacy of the therapeutic delivery platform^29^. Furthermore, since Wnt7a pre-transplant treatment improves MuSC engraftment efficiency and donor fiber dispersion^73^, we speculate that we might expect an additive improvements if the Wnt7a pre-treated therapeutic cell population was delivered in an optimized delivery scaffold.

Together, our study aimed at optimizing the MuSC delivery method and exploring the mechanistic underpinnings of our in vivo results pave the way for reducing the number of cells required for cell transplantations to make MuSC transplant therapy more feasible and cost-effective.

## Acknowledgements

We would like to thank Ms. Aymin Mumtaz for her technical assistance in these studies. This study was funded by the Natural Sciences and Engineering Research Council (CREATE ToEP fellowship to S.D., RGPIN 435724-13 and Canada Research Chair 950-231201 to P.M.G.); Ontario Provincial Government (ER15-11-073 to P.M.G.); Canada Foundation for Innovation (31390 to P.M.G.); Toronto Western Arthritis Program (to P.M.G.); University of Toronto Faculty of Medicine Dean’s Fund (to P.M.G.); and Canadian Institutes of Health Research (ONM-137370 to P.M.G.).

## Materials & Methods

### Animals

All animal use protocols were reviewed and approved by the Division of Comparative Medicine (DCM) at University of Toronto. C57Bl6/NCrL (Charles River) and C57BL/6-Tg(CAG-EGFP)1Osb/J (expressing eGFP under the control of Actin, Jackson Laboratories), B6.129(Cg)-Cd44tm1Hbg/J (CD44^−/−^ mice courtesy of Dr. Tak Mak, University of Toronto^74^), Pax7-zsGreen reporter mice (courtesy of Dr. Michael Kyba, University of Minnesota^75^) were used in this project. The C57BL/6-Tg(CAG-EGFP)1Osb/J and Pax7-zsGreen lines were maintained as a heterozygous line by breeding against wild-type C57Bl6/NCrL females, resulting in litters comprised of both wild type and transgenic pups. The B6.129(Cg)-Cd44tm1Hbg/J line was maintained as a homozygous line. 8-12 week old mice (wild type or transgenic) were used in all of the experiments. Injuries to the *tibialis anterior* (TA) muscle were induced by injecting 30 µL of a 2% BaCl_2_ (Bio Basic, cat. no. BC2020) into the center of the TA muscle of anaesthetized mice using a 100 µL insulin syringe (BD, cat. no. 324702).

### Hyaluronan (HA) and methylcellulose (MC) preparation

Sodium Hyaluronate (HA; Kibun Food Chemifa, cat. no. 9004-61-9) and methylcellulose (MC; Shin Etsu, cat. no. SM-4000) were dissolved in distilled H_2_O at 0.5 and 1 g / L at 4°C overnight. HA and MC solutions were then sterile filtered using a 0.2 µm vacuum filter and aliquoted in 35 mL aliquots in 50 mL conical tubes and flash frozen in liquid N_2_. The caps were then replaced with perforated, filter caps (Corning, cat. no. 431720) and placed in −80°C for 2 hours. The tubes were then lyophilized for 2 - 3 days. The resulting sterile HA and MC were then weighed out at a 1:1 ratio, depending on the amount of HAMC required, and dissolved in the appropriate solution at a 0.75:0.75 % weight/volume ratio overnight at 4°C on a rocking shaker.

### Primary myoblast isolation

Primary myoblast lines were established using a method similar to the one described in Rando et al^60^. Briefly, all of the hind limb muscles were dissected from humanely euthanized 8-12 week old mice and placed in 7 mL DMEM containing Type IA collagenase (Sigma, cat. no. C9891) at 628 Units / mL. The muscles were then dissociated using a gentleMACS Dissociator (Miltenyi Biotec, cat. no. 130-093-235). Tissues were then incubated on a rocking shaker at 37°C and 5% CO_2_ for 90 minutes. Dispase II (Life Technologies, cat. no. 17105041) was added at a concentration of 4.8 U/mL and the muscle was incubated for another 30 minutes on the rocking shaker at 37°C and 5% CO_2_. After the incubation, the tissues were passed through a 20 G needle equipped with a 10 mL syringe to fully dissociate the tissue. 7 mL of growth media was then added to the dissociated tissue and the entire volume was filtered using a 40 µm cell strainer (Corning, cat. no. 352340). The filtered liquid was centrifuged at 400g for 15 minutes. The supernatant was then discarded and the pellet resuspended in 1 mL of red blood cell lysis buffer (0.155 M NH_4_Cl, 0.01 M KHCO_3_, 0.1 mM EDTA), which was then incubated at room temperature for 7 minutes. 10 mL of FACS buffer (PBS, 2.5% goat serum, 2 mM EDTA) was added and centrifuged at 400 G for 15 minutes. The pellet was then resuspended in culture media and transferred to a collagen coated 10 cm tissue culture plate. The culture media was changed the following day. After 2 days, the tissue culture plate was washed once with PBS and 5 mL of PBS is added to the tissue culture plate. The plate is placed at 37°C and 5% CO_2_ for ~7-8 minutes after which the mitotic satellite cells are detached from the plate by firmly tapping the plate. The PBS containing the cells is collected, centrifuged, and the cells are transferred in culture media to a new collagen-coated tissue culture plate. This process is repeated every few days until the majority of the cells are myoblasts.

### Muscle stem cell isolation

Muscle stem cells were obtained using a similar method described in Sacco et al^13^. Briefly, all of the hind limb muscles were dissected from humanely euthanized 8-12 week old mice and placed in 7 mL DMEM containing Type IA collagenase at 628 Units/mL. The muscles were then dissociated using a gentleMACS Dissociator. Tissues were then incubated on a rocking shaker at 37°C and 5% CO_2_ for 90 minutes. Dispase II was then added at a concentration of 4.8 U / mL and the muscle was incubated for another 30 minutes on the rocking shaker at 37°C and 5% CO_2_. After the incubation, the tissues were passed through a 20 G syringe needle equipped with a 10 mL syringe to fully dissociate the tissue. 7 mL of serum containing media (10% FBS, 90% DMEM) was then added to the dissociated tissue and the entire volume was filtered using a 40 µm cell strainer. The filtered liquid was centrifuged at 400g for 15 minutes. The supernatant was then discarded and the pellet, resuspended in 1 mL of red blood cell lysis buffer, was incubated at room temperature for 7 minutes. 10 mL of FACS buffer was added and the cell slurry centrifuged at 400 G for 15 minutes. The pellet was then resuspended in 1 mL of FACS buffer. The cells were incubated with biotinylated CD31 (1:200, BD, cat. no. 553371), CD45 (1:500, BD, cat. no. 553078), CD11b (1:200, BD, cat. no. 553309), and Sca1 (1:200, BD, cat. no. 553334) for 30 minutes on a rocking shaker at 4°C. After washing the cells and resuspending them in 1 mL FACS buffer, streptavidin microbeads (1:20, Miltenyi Biotec, cat. no. 130-048-101), streptavidin PE-Cy7 (1:200, Life Technologies, cat. no. SA1012), α7 Integrin – PE conjugated (1:500, AbLab, cat. no. 530010-05), and CD34-eFluor 660 conjugated (1:65, ebioscience, cat. no. 50-0341-82) antibodies were added to the cells. The cells were incubated for 1 hour on the rocking shaker at 4°C. After the incubation, the cells were once more washed with FACS buffer, and depleted for biotin labeled cells by passing through a magnetic column (Miltenyi Biotec, cat. no. 130-091-051) after resuspension. The cells were washed one final time and resuspended in 1 mL FACS buffer containing Propidium Iodide (Sigma, cat. no. P4864). Muscle stem cells (MuSCs) were then sorted using a BD FACS-Aria II based on 7-AAD^−^/CD31^−^/CD45^−^/CD11b^−^/Sca1^−^/CD34^+^/α7-Integrin^+^.

### Myogenic cell culture

Muscle stem cells (MuSCs) and primary myoblasts (pMBs) were cultured in tissue culture plates coated with Type 1 rat tail Collagen protein (Life Technologies, cat. no. A1048301). The culture medium was composed of 20% fetal bovine serum (Life Technologies, cat. no. 12483), 79% F-12, 1% Penicillin-Streptomycin (Life Technologies, cat. no. 11765054) and rh-bFGF (ImmunoTools, cat. no. 11343627) at a final concentration of 2.5 ng / mL. The culture media was changed every other day and the cells were maintained at 37°C and 5% CO_2_.

### Myogenic cell transplantation

The tibialis anterior muscle of 8-12 week old wild type offspring of crossing C57Bl6/N and actin-eGFP transgenic mice were injured with a single intramuscular injection of BaCl_2_ (described in animal section). Two days post-injury, muscle stem cells were freshly isolated from the transgenic siblings of the injured mice. The freshly isolated muscle stem cells were then suspended in either saline or 0.75:0.75 HA:MC dissolved in saline.

The injured wild type mice were then anaesthetized in the operating room. The hair was removed from the skin on top of the TA muscle and a small 1” incision was created in the skin to expose the TA muscle. 2 µL of saline or HAMC containing 1.5×10^3^, 5×10^3^, or 10×10^3^ cells was injected into the center of the exposed muscle using a 32G needle and a Hamilton syringe. The skin was then closed and sutured using a 6-0 suture (Covidien, cat. no. SS681). The procedure was performed on both TA muscles of the mice, with one muscle receiving the cells delivered in saline, and the other, cells delivered within HAMC.

3-4 weeks post-transplantation, the mice were euthanized and the TA muscles were harvested and either fixed and frozen for immunohistochemistry or dissociated for flow cytometry analysis.

Similar methods were used to transplant myoblasts, HAMC, and saline into the muscle.

### Immunohistochemistry

Post-euthanasia, the TA muscles were extracted and fixed in 0.5% Pierce methanol-free paraformaldehyde (Thermo Fisher, cat. no. 28908) for 2 hours at 4° C. The muscles were then transferred to a 20% sucrose (Sigma, cat. no. S9378) solution in PBS overnight at 4° C. They were then embedded and frozen in OCT (TissueTek, cat. no. 4583) and stored at −80° C prior to further processing.

Frozen tissues were sectioned and mounted as 10 µm sections. The tissue sections were rehydrated, blocked, and permeabilized with blocking solution consisting of 20% goat serum (Life Technologies, cat. no. 16210-072), 79% PBS, and 1% Triton-X100 (Bioshop, cat. no. TRX777) for 1 hour at room temperature (RT) or overnight at 4°C. 1° antibody was added to the slides and incubated for 2 hours at RT or overnight at 4°C. After numerous washes with PBS, 2° antibody solutions were added to the slides and incubated for 30 minutes at RT. The slides were then mounted using fluoromount (Sigma, cat. no. F4680) and subsequently imaged.

The primary antibodies used in this project were rabbit anti-GFP (1:500, Invitrogen, cat. no. A11122), rat anti-CD68 (1:200, abcam, ab53444), and rat anti-Ly6G (1:500, abcam, ab25377). Secondary staining was performed using Alexafluor goat anti-rabbit 488 (1:500, Life Technologies, cat. no. A11008), Alexafluor goat anti-rat 647 (1:500, Life Technologies, cat. no. A21247). The nuclei were visualized using Hoechst 33342 (1:1000, Life Technologies, cat. no. H3570).

### Flow cytometry Analysis

Flow cytometry analysis was used to enumerate transplanted GFP^+^ myoblasts 2, 24, and 48 hours post-transplantation into the injured TA. Post-euthanasia, the TA muscles were extracted and placed in 1 mL DMEM containing Type IA collagenase at 628 Units/mL and dispase II at 4.8 U/mL. The muscle was minced with scissors into small pieces (roughly 2 mm × 2 mm × 2 mm pieces) and then placed on a rocking shaker at 37°C and 5% CO_2_ for 120 minutes. After the incubation, the tissues were passed through a 20 G syringe needle equipped with a 10 mL syringe to fully dissociate the tissue. The entire volume was then transferred to a FACS tube equipped with a cell strainer cap (Corning, cat. no. 352235) to further deplete debris. 3 mL of FACS buffer was then added to the sample, which was then centrifuged at 400 g for 15 minutes. The supernatant was discarded and the pellet was resuspended in 0.5 mL FACS buffer. Propidium iodide was added at 1:1000. The sample was then analyzed on a BD-Canto flow cytometer (courtesy of Dr. Peter Zandstra, University of Toronto), to enumerate PI^−^/GFP^+^ cells.

For CD44 expression analysis, a method similar to muscle stem cell isolation protocol was followed. In this case, the cell slurry was incubated with biotinylated CD31, CD45, CD11b, and Sca1 for 60 minutes on the rocking shaker at 4°C. Either rat anti-mouse CD44 (1:100, BD, cat. no. 558739) or purified rat IgG1, κ Isotype Control (1:100, BD, cat. no. 559072) was also added in this step. After washing the cells and resuspending them in 1 mL FACS buffer, Streptavidin PE-Cy7, α7 Integrin – PE conjugated, and CD34-FITC conjugated (1:65, ebioscience, cat. no. 11-0341-82), and Alexafluor goat anti-rat AF647 antibodies were added to the cells. The cells were incubated for 1 hour on the rocking shaker at 4°C. The cells were washed one final time and resuspended in 1 mL FACS buffer containing propidium iodide (1:1000). The percentage of healthy or activated MuSCs expressing CD44 was determined using a BD FACS-Aria II flow cytometer based on 7-AAD^−^/CD31^−^/CD45^−^/CD11b^−^/Sca1^−^/CD34^+^/α7-integrin^+^/CD44^+^.

### EdU Analysis

Freshly isolated MuSCs were seeded into 96 well plate tissue culture plates in either 0.75:0.75 HAMC or culture media. Cells that had entered cell cycle were visualized using the Click-it EdU kit (Thermo Scientific, cat. no. C10634). The cells were pulsed with 5-ethynyl-2’-deoxyuridine (EdU) for 12 hours, after which they were trypsinized and cytospun (courtesy of Dr. Julie Audet, University of Toronto) onto charged glass slides. The cells were immediately fixed with a 4% paraformaldehyde solution in ddH_2_O. The EdU was then visualized using Alexa Fluor 647 azide, and the nuclei were counter-stained with Hoechst 33342.

### Cell loss, ejection, and viability assays

Passage 4-10 primary myoblasts were seeded into 96 well plate tissue culture plate in either 0.75:0.75 HAMC or culture media. The Live/Dead^®^ assay (Life Technologies, cat. no. L3224) was used to determine cell viability at desired time points. Calcein AM (1:2000), Ethd-1°(1:500), and Hoechst 33342 (1:1000) were used to determine live, dead, and nuclei respectively. The cells were incubated at 37°C and 5% CO_2_ for 15 minutes with the live/dead stain prior to imaging.

The viability of the ejected MuSCs and pMBs was assessed by ejecting 10 µL of the cells using 0.75:0.75 HA:MC or growth media into a collagen coated 384 well plate. The Live/Dead^®^ assay was then used at the desired time points post-ejection to determine cell viability, as described previously.

Ejection efficiency assays were performed by ejecting cells suspended in 0.75:0.75 HA:MC or growth media at 500,000 cells/mL using a 29 G syringe onto a hemocytometer and counting the number of ejected cells.

Injured tissue extract was obtained by lysing the injured TAs of the mice in F-12 media. The lysis’ protein content was determined using Pierce BCA Protein Assay kit (Thermofisher, 23227). The cells were cultured with a final concentration of 1 mg/ml of the extract.

### Dispersion analysis

Dispersion of the GFP^+^ fibers was determined by calculating the average of the average distance of each GFP^+^ fiber to all other GFP^+^ fibers. A higher average would mean the fibers are more dispersed and spread out throughout the tissue. ImageJ analysis and Matlab coding were used to determine dispersion using the following formula:

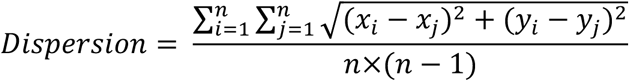
where
n is the number of GFP^+^ fibers *x_i_* and *y_i_* are the coordinates of the center of the GFP^+^ fibers.

### Imaging and microscope

Tissue sections and cells were visualized using an Olympus IX83 inverted microscope and 10x or 20x objectives. The images were captured with an Olympus DP80 dual CCD color and monochrome camera and CellSens software. Images were adjusted consistently using open source ImageJ software.

### Statistical analysis

All of the experiments were performed with a minimum of three biological replicates. Paired student’s t-test was used in all statistical analysis, unless otherwise specified, with significance set at p < 0.05.

---

**Supplemental Figure 1.**
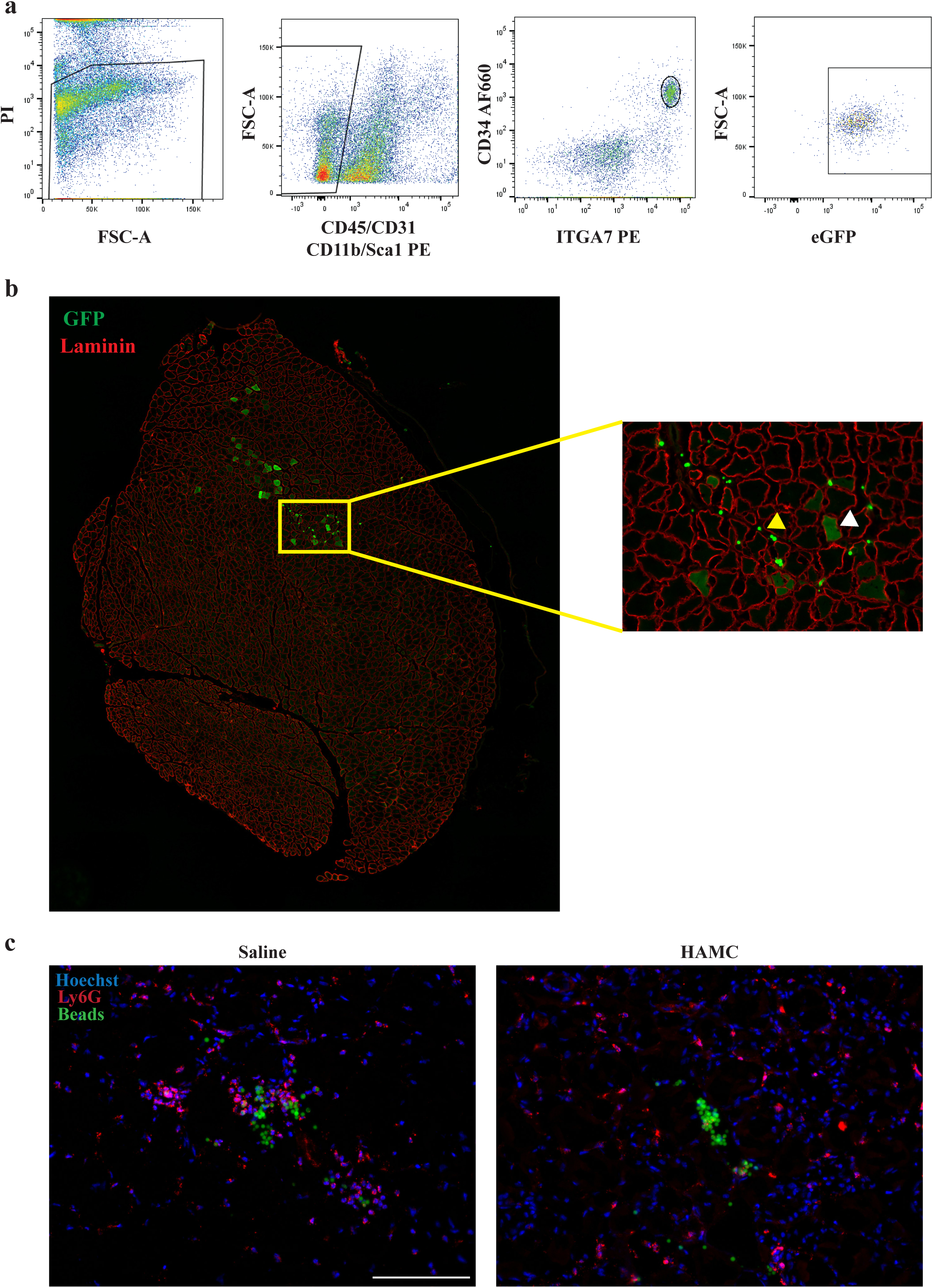
Muscle stem cell sorting and injection methods. **(a)** Representative fluorescence activated cell sorting (FACS) gates for isolation of live (far left plot, PI^−^), lineage negative (middle left plot, CD31^−^/CD11b^−^/CD45^−^/Scal^−^), CD34^+^/ITGA7^+^ (middle right plot) MuSCs from GFP^+^ (far right plot) mice. **(b)** Representative tiled confocal image of a transverse section from a *tibialis anterior* muscle that was harvested and immunostained 4 weeks after transplantation with GFP^+^ (green) MuSCs. Transplant location was determined by the presence of co-transplanted fluorescent (GFP^+^) spherical microbeads (yellow arrowheads) that are readily visible in the blown-up image (right). Donor-derived GFP^+^ fibers (white arrowheads) are viewed in the vicinity as well as distant locations from the transplantation location. **(c)** Representative epifluorescence images of fluorescent bead distribution one-month following intramuscular transplant into BaCl2 pre-injured TA muscle when delivered within saline (left) compared to HAMC (right). Tissue sections are co-stained to visualize neutrophil (Ly6G, red) and nuclei (Hoechst, blue). Scale bar, 100 µm.

**Supplemental Figure 2.**
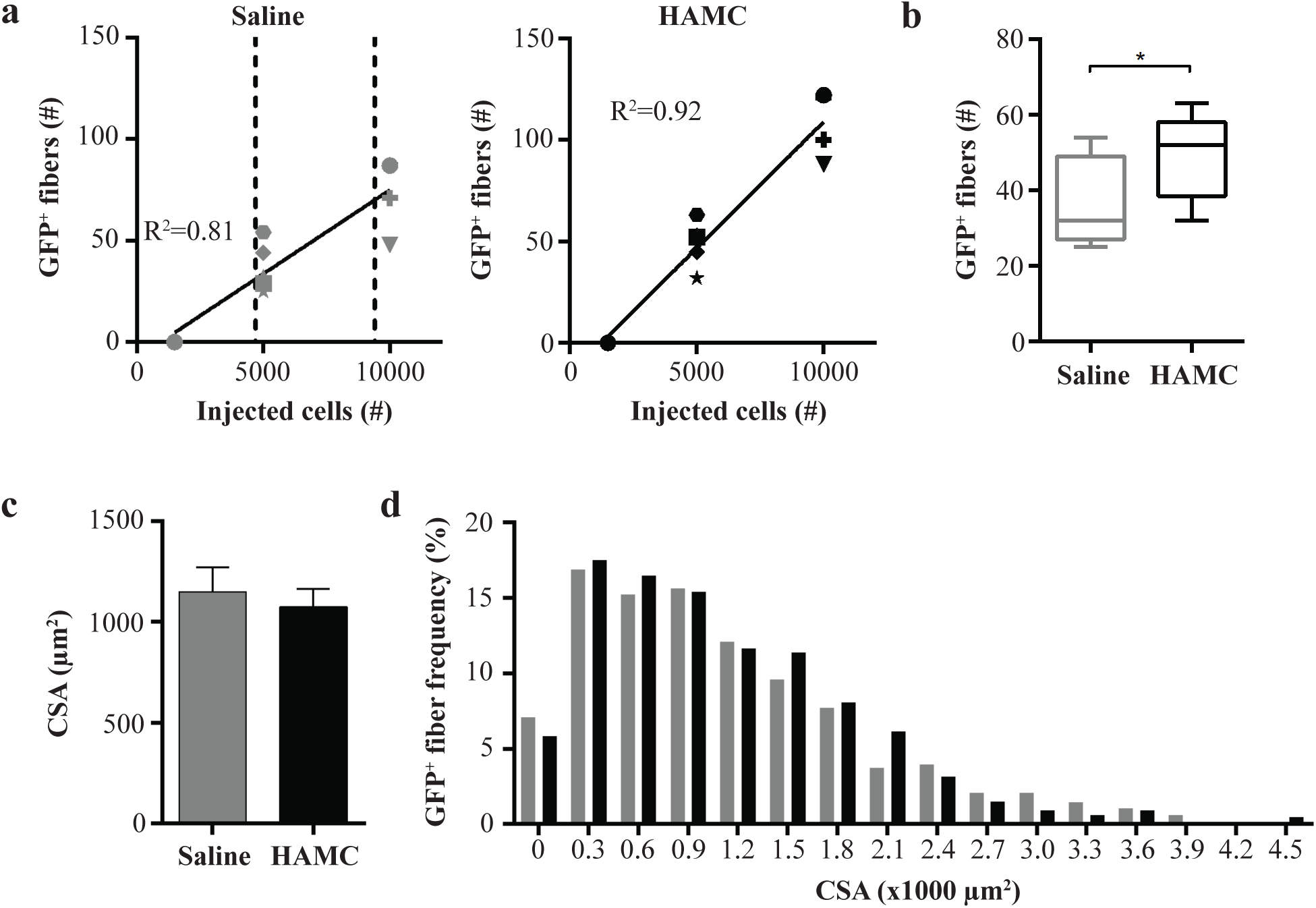
Analysis of MuSC engraftment in immunocompetent hosts. **(a)** Paired scatter plots comparing the number GFP^+^ fibers quantified in recipient animals one-month following injection of 1.5×10^3^ (n = 2), 5×10^3^ (n = 5), and 10×10^3^ (n = 4) freshly isolated GFP^+^ MuSCs that were delivered intramuscularly within **(left)** saline or **(right)** HAMC. Please note that the data in these paired scatter plots were used to generate the Figure 1c and Supplemental Figure 2b bar graphs. **(b)** Bar graph indicating the number of GFP^+^ fibers one-month following intramuscular injection of 5×10^3^ MuSCs within saline or HAMC. n = 5. **(c)** Bar graph showing the average cross-sectional area (CSA) of GFP^+^ donor-derived fibers arising from GFP^+^ MuSCs delivered intramuscularly within saline (dark grey) or HAMC (black). n = 9. **(d)** Histogram displaying donor-derived GFP^+^ fiber CSA comparing saline (dark grey) to HAMC (black) delivery of GFP^+^ MuSCs. Statistical significance determined by student’s t-test where; p < 0.05.

**Supplemental Figure 3.**
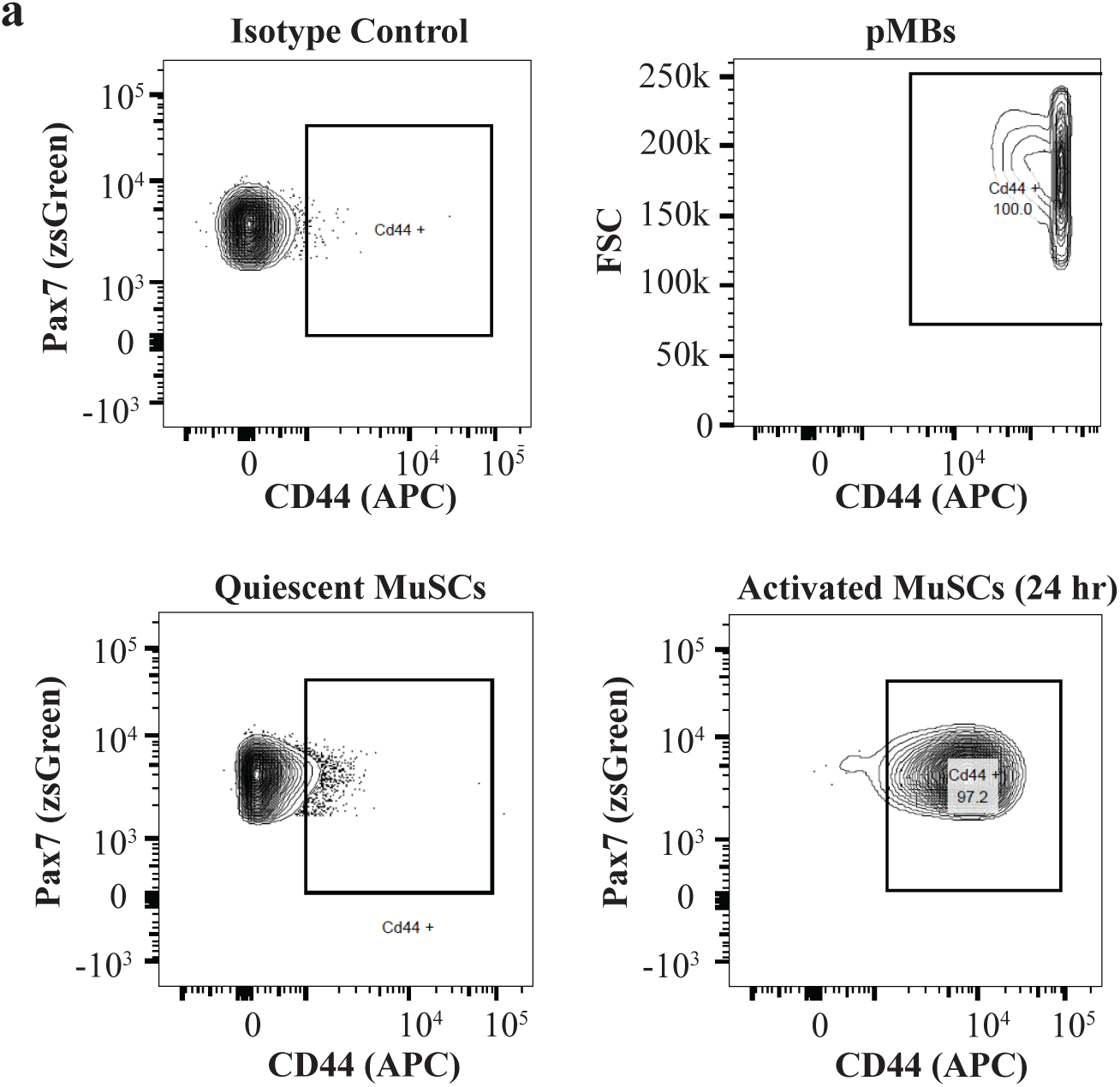
CD44 is differentially expressed within the myogenic hierarchy. **(a)** Representative flow cytometric plots analyzing CD44 cell surface expression on freshly isolated quiescent (bottom left) and activated (bottom right; 24 hours post-BaCl_2_ injury) MuSCs, and primary myoblasts (top right), as compared to isotype control staining (top left).

**Supplemental Figure 4.**
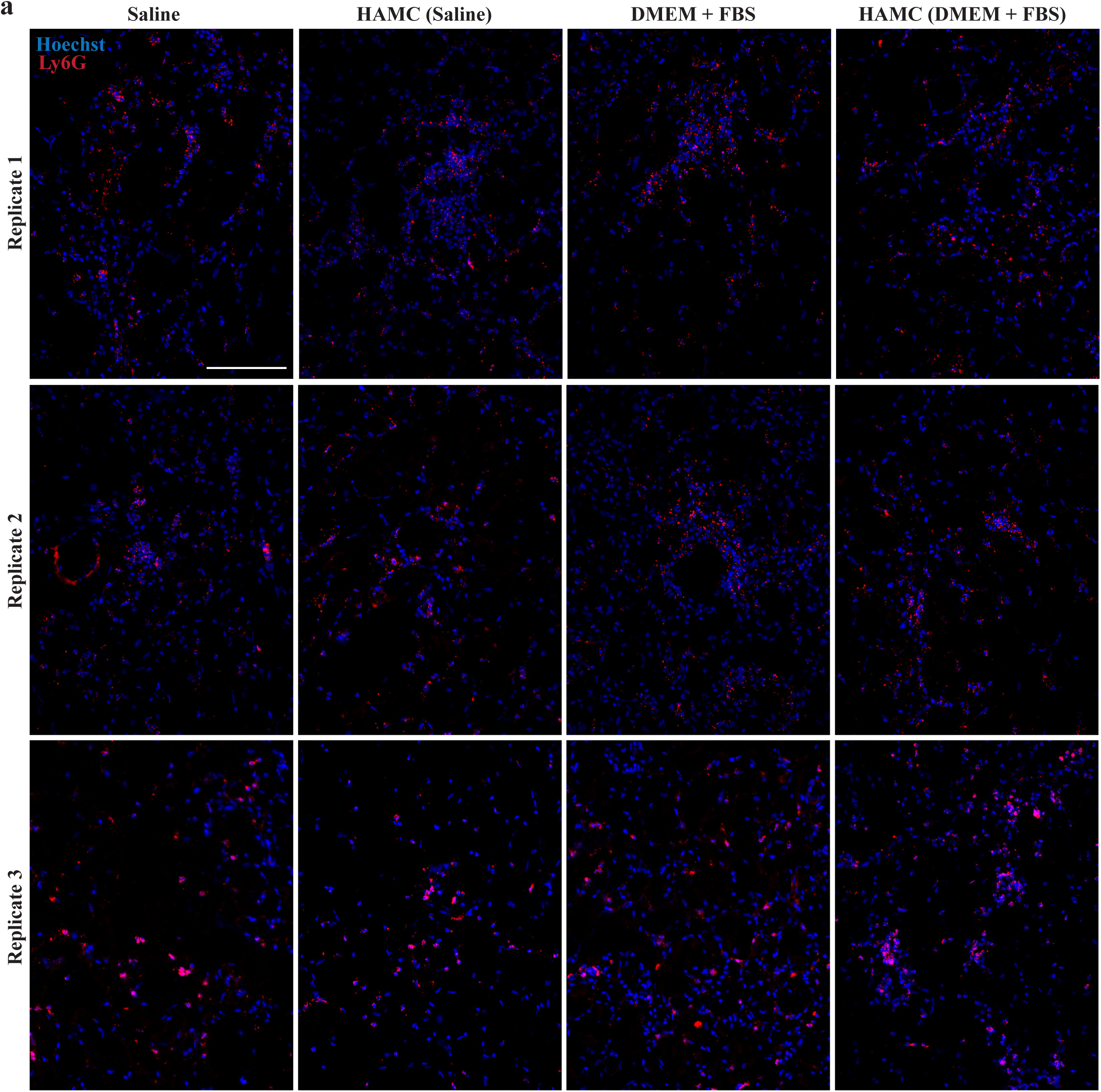
HAMC does not alter the presence of Ly6G^+^ cells in regenerating skeletal muscle at early time-points post-injection. **(a)** Transverse sections of skeletal muscle tissue immunostained with Ly6G (red) to visualize neutrophils and counterstained with the nuclear stain Hoechst (blue). Shown are representative epifluorescence images from each of three replicate animals (rows) that were injured with a BaCl2 injection and then 2-days later were injected with saline (far left column), HAMC reconstituted in saline (middle left column), serum containing media (middle right column), or HAMC reconstituted in serum containing media (far right column) and sacrificed for analysis 2 hours later. n=3. Scale bar, 100 µm.

**Supplemental Figure 5.**
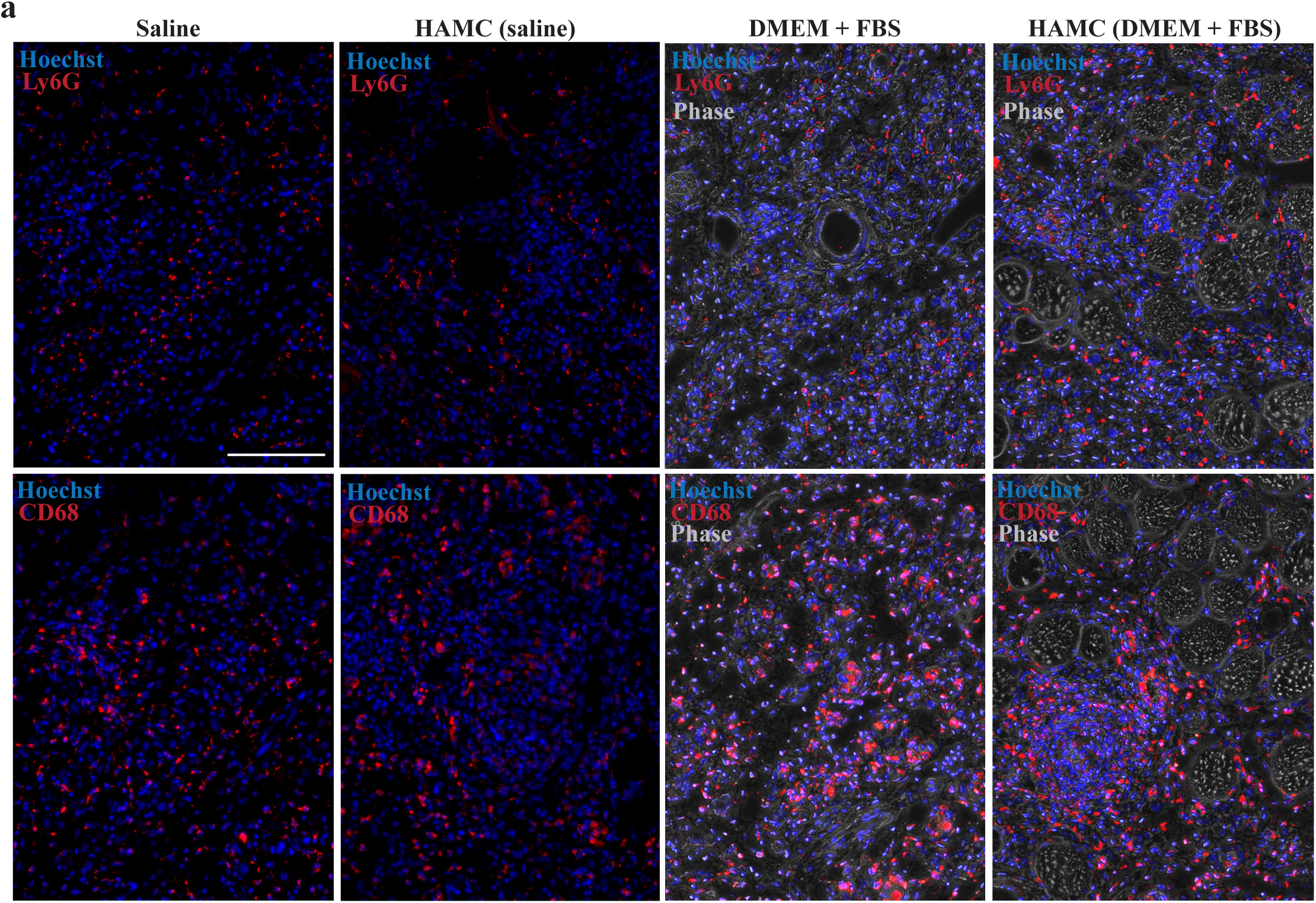
HAMC does not alter the presence of Ly6G^+^ or CD68^+^ cells in regenerating skeletal muscle at later time-points post-injection. **(a)** Transverse sections of skeletal muscle tissue immunostained with Ly6G (red; top row) to visualize neutrophils, or CD68 (red, bottom row) to visualize macrophages, and counterstained with the nuclear stain Hoechst (blue). Shown are representative epifluorescence images from animals that were injured with a BaCl2 injection and then 2-days later were injected with saline (far left column), HAMC reconstituted in saline (middle left column), serum containing media (middle right column), or HAMC reconstituted in serum containing media (far right column) and sacrificed for analysis 24 hours later. Scale bar, 100 µm.

